# Testing the Tea Bag Index as a potential indicator for assessing litter decomposition in aquatic ecosystems

**DOI:** 10.1101/2021.04.26.441560

**Authors:** Taiki Mori, Kenji Ono, Yoshimi Sakai

## Abstract

The Tea Bag Index (TBI) approach is a standardized method for assessing litter decomposition in terrestrial ecosystems. This method allows determination of the stabilized portion of the hydrolysable fraction during the decomposition process, and derivation of a decomposition constant (*k*) using single measurements of the mass-loss ratios of green and rooibos teas. Although this method is being applied to aquatic systems, it has not been validated in these environments, where initial leaching tends to be higher than in terrestrial ecosystems. Here, we first validated a critical assumption of the TBI method that green tea decomposition plateaus during the standard incubation period of 90 days, and then tested the accuracy of a TBI-based asymptote model using a second model obtained from fitting actual decomposition data. Validation data were obtained by incubating tea bags in water samples taken from a stream, a pond, and the ocean in Kumamoto, Japan. We found that green tea decomposition did not plateau during the 90-day period, contradicting a key assumption of the TBI method. Moreover, the TBI-based asymptote models disagreed with actual decomposition data. Subtracting the leachable fraction from the initial tea mass improved the TBI-based model, but discrepancies with the actual decomposition data remained. Thus, we conclude that the TBI approach, which was developed for a terrestrial environment, is not appropriate for aquatic ecosystems. However, the use of tea bags as a standard material in assessments of aquatic litter decomposition remains beneficial.

## Introduction

Litter decomposition is an important process in aquatic systems (Graça et al., 2015), playing a major role in the global cycling of carbon and nutrients (Battin et al., 2009; Gessner et al., 1999; Graça et al., 2015). Three main factors control aquatic litter decomposition: litter quality (Jabiol et al., 2019; Neiff et al., 2006); environmental factors such as temperature, nutrient availability, salinity, acidity, and oxygen concentration (Almeida Júnior et al., 2020; Ferreira et al., 2015; Gomes et al., 2018; Griffiths and Tiegs, 2016; Woodward et al., 2012; Young et al., 2008); and decomposer community composition, including macroinvertebrates, fungi, and bacteria (Balibrea et al., 2020; Hieber and Gessner, 2002).

Litter bags filled with litter from the local area are often used to assess litter decomposition rates in aquatic systems. However, understanding the impacts of environmental changes on litter decomposition rates at large geographical scales requires a standardized method (Keuskamp et al., 2013; Mori et al., 2021a), because high local variation in litter quality may negate the influence of environmental effects. A standardized method applicable to both aquatic and terrestrial ecosystems would be beneficial (Seelen et al., 2019), because integrated models of both terrestrial and aquatic systems remain poor (García-Palacios et al., 2016).

The Tea Bag Index (TBI) approach proposed by Keuskamp et al. (2013) is a standardized method for assessing litter decomposition that was developed in a terrestrial ecosystem. This method uses two types of commercially available tea bags (green and rooibos teas, Lipton) as standard materials to calculate the TBI, which consists of two parameters: a stabilization factor *S* (the stabilized portion of the hydrolysable fraction during decomposition) and the decomposition constant *k* of an asymptote model (Keuskamp et al., 2013). Due to its cost-effectiveness and ability to collect comparable globally distributed data (Keuskamp et al., 2013), multiple studies have used the TBI (Becker and Kuzyakov, 2018; Fanin et al., 2020; Fujii et al., 2017; Mori et al., 2021b; Mueller et al., 2018; Petraglia et al., 2019).

Calculation of the TBI rests on several assumptions. *S* is calculated as the ratio between the mass of hydrolysable fraction of green tea remaining at the end of the 90-day standard incubation period and the entire hydrolysable fraction of green tea (0.842; Keuskamp et al. 2013); this assumes that the decomposition of the acid-insoluble fraction of green tea is negligible, and that the undecomposed hydrolysable fraction stabilizes and transforms to a recalcitrant fraction within the incubation period. Thus, the decomposition curve of green tea must reach a plateau before the end of the incubation period, given that the decomposition and stabilization of the hydrolysable fraction should be completed within this period. Next, the decomposition constant (*k*) is determined by fitting an asymptote model to rooibos tea decomposition data using the following formula:

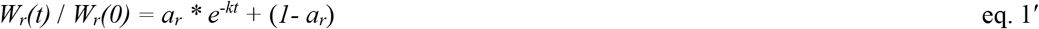

where *W*_*r*_*(t)* is the ratio of the remaining mass of rooibos tea after the incubation period *t* relative to the initial mass, *k* is a decomposition constant, *a*_*r*_ is the decomposable fraction, and *1- a*_*r*_ is the undecomposable fraction of rooibos tea (Keuskamp et al., 2013). The TBI approach allows parameterization of the asymptote model without fitting the model to time series data of rooibos tea decomposition by determining *a*_*r*_ using green tea decomposition data. Because the TBI approach assumes that the acid-insoluble fraction does not degrade within a 90-day period, *a*_*r*_ represents the decomposable hydrolysable fraction (1 – acid-insoluble fraction) of rooibos tea. Under the assumption that the ratio of the decomposable hydrolysable fraction to the total hydrolysable fraction of rooibos tea is the same as that of green tea, *a*_*r*_ can be calculated as the total hydrolysable fraction (0.552; Keuskamp et al. 2013) multiplied by (1 –*S*). The decomposition constant *k* can be determined by substituting the values obtained from a single measurement of rooibos tea decomposition for the parameters in eq. 1 (i.e., *t* is 90 and *W*_*r*_*(t)* is the remaining-mass ratio of rooibos tea at the end of the incubation period).

Although the TBI method was developed in a terrestrial ecosystem, it has increasingly been adopted in aquatic environments (Hunter et al., 2019), including mangrove systems (Mueller et al., 2018), marshes (Mueller et al., 2018; Yousefi Lalimi et al., 2018), and streams (Peralta-Maraver et al., 2019). In these studies, the TBI was calculated following the protocol of Keuskamp et al. (2013), which assumes that the index can be applied equivalently in terrestrial and aquatic ecosystems. However, the initial decomposition phase, i.e., leaching, is much more rapid in aquatic environments (Webster and Benfield, 1986), which could cause the TBI-based asymptote model to deviate from real-world decomposition (Seelen et al., 2019). In addition, the assumption that green tea decomposition plateaus during the incubation period has not been verified in an aquatic ecosystem (Keuskamp et al., 2013).

We first aimed to validate whether green tea decomposition does plateau during the 90-day incubation period, and then tested the accuracy of a TBI-based asymptote model by comparison with a model derived from actual decomposition data. Seelen et al. (2019) suggested that correcting the initial tea weights by subtracting the easily leachable fraction may provide a more accurate TBI in aquatic systems. However, the validity of this corrected approach was not evaluated using actual data. Therefore, we also evaluated a TBI-based model corrected by subtraction of the easily leachable fraction.

## Materials and methods

### Incubation experiment

We evaluated application of the TBI approach to aquatic ecosystems by monitoring the mass-loss ratios of green tea bags submerged in water. A microcosm approach (Santschi et al., 2018), rather than an *in-situ* approach, was used to control environmental variability as much as possible. We collected water samples for our experiment from three different sources: a stream running through an evergreen conifer plantation dominated by *Cryptomeria japonica* (Linnaeus f.) D. Don and *Chamaecyparis obtusa* (Sieb. et Zucc.) Endl. in Yamaga City, an artificial pond located at the Forestry and Forest Products Research Institute (FFPRI) in Kyushu, and the ocean in Uki City. All three sites are located in Kumamoto Prefecture, Japan. Because the purpose of our study did not include investigating factors leading to differences in decomposition rates among the water samples, we did not perform chemical analyses of the water samples. Each of the three water samples was placed into eight plastic bottles (6.5 cm in diameter); four were assigned to green tea (EAN: 87 22700 05552 5, non-woven mesh, Lipton) and four to rooibos tea (EAN: 87 22700 18843 8, non-woven mesh, Lipton). We placed 200 mL of water in each bottle and submerged five tea bags therein. We then covered the bottles with a polyethylene sheet to prevent evaporation (Mori et al., 2013). The bottles were incubated at 25°C in the dark. One tea bag per replicate was retrieved at 3, 11, 27, 55, and 91 days after the start of the incubation period, whereupon weights were determined by oven drying at 70°C for 72 h.

### Calculating TBI

The TBI was calculated following Keuskamp et al. (2013). *S* was calculated as:

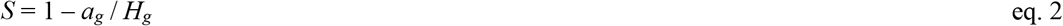

where *a*_*g*_ is the mass loss of green tea during the 90-day period (note that we used 91 days) and *H*_*g*_ (0.842) is the hydrolysable fraction of green tea (Keuskamp et al. 2013). Assuming that the *S* value of rooibos tea is the same as that of green tea, the decomposable fraction of rooibos tea was calculated as:

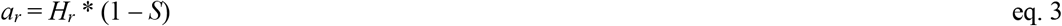

where *H*_*r*_ (0.552) is the hydrolysable fraction of rooibos tea (Keuskamp et al. 2013). The decomposition constant (*k*) was determined from an asymptote model of rooibos tea decomposition:

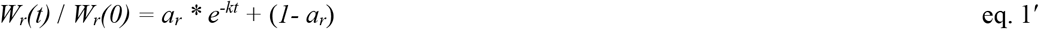

where *W*_*r*_*(t)* and *W*_*r*_*(0)* are the mass of rooibos tea remaining after the incubation time and that before the incubation time *t*, respectively.

### Leaching factor correction

Seelen et al. (2019) proposed correction of the TBI using a leaching factor, which was defined as the ratio of the easily leachable fraction to the total tea weight. Initial tea weight can then be corrected by multiplying the initial tea mass by (1 – the leaching factor). Accordingly, all remaining mass values and hydrolysable fractions of both the green and rooibos teas (*H*_*g*_ and *H*_*r*_) were corrected by subtracting the leaching factor and dividing the result by (1 – the leaching factor). The TBI and the TBI-based asymptote model were then calculated using the corrected data.

Seelen et al. (2019) suggested that a leaching factor be determined using mass loss data of both green and rooibos teas after submersion in water for 3 h. We chose to calculate a corrected TBI using several scenarios with different combinations of leaching factors for green and rooibos teas, rather than experimentally determining leaching loss in the manner suggested. We chose this approach because it has been reported that leaching can occur for up to 72 and 48 hours in green and rooibos teas, respectively (Edwartz, 2018), and a long-term leaching experiment could overestimate the leaching factor due to microbial decomposition. In addition, solute concentration may influence initial leaching losses in tea, leading to differences between ecosystems and potential interactions with the teas’ initial chemical composition. If the TBI is to be corrected by a leaching factor, then a global leaching factor would be most appropriate. Our approach therefore represents the first step toward a global leaching factor applicable to all aquatic environments. We created multiple scenarios, with a maximum leaching factor based on the mass loss of the teas on day 3 of incubation, assuming that leaching would be complete within 3 days and that the minimum leaching factor is 0, and intermediate values between the maximum leaching factor and zero, in our calculations of a corrected TBI. Asymptote models were constructed using the corrected TBI. We note that if a leaching factor is applied in assessments of aquatic litter decomposition, the results will not be comparable to those in terrestrial systems due to inherent differences in initial chemical compositions.

### Leaching experiments

We assessed tea leaching in deionized water and a tea solution to understand the influence of solute concentration on leaching. We used a tea solution rather than a salt solution to avoid any potential chemical reactions that could influence the leaching process. Solutions of both green and rooibos teas were prepared by submerging 10 tea bags in 600 mL of deionized water overnight. The deionized water and tea solutions were autoclaved at 120°C for 20 min for sterilization purposes. The autoclaved solutions were then placed in an incubator overnight at 25°C. Green and rooibos tea bags were then submerged into either the deionized water or tea solution, where green and rooibos tea bags were submerged in green and rooibos solutions, respectively, for 20 min or 9 h. The dry weights of the tea bags were then determined following oven drying at 70°C for 72 h.

### TBI-based asymptote model in terrestrial systems

As a reference, we calculated a decomposition curve based on a TBI-based asymptote model in a terrestrial ecosystem using published data (Keuskamp et al., 2013). These data, representing the outcomes of a laboratory incubation study conducted at 25°C, were obtained from Keuskamp et al. (2013) using the Data Thief 3.0 program (Tummers, 2006). We compared the TBI-based model to an asymptote model (eq. 1) that had been fitted to the time series data. Given that Keuskamp et al. (2013) did not collect remaining-mass observations on day 90 of incubation, the remaining mass of both green and rooibos teas at 90 days was calculated using the fitted asymptote model with a *t* of 90.

### Statistical analyses

All statistical analyses were performed using R software (R Core Team, 2019). The asymptote model was fitted using nonlinear regression. Additionally, we fitted a double exponential model to the experimental data obtained from the stream- and pond-water samples using nonlinear regression. The model was expressed as:

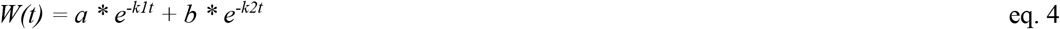

where *W(t)* is the mass remaining after *t, k1* and *k2* are decomposition constants, and *a* and b represent organic matter fractions differing in decomposability. A one-way ANOVA followed by Tukey’s post hoc test was used to compare the remaining-mass ratios of the teas among the different water samples. Shapiro-Wilk tests showed that the data were not significantly different from a normal distribution, and log-transformation did not improve data fit; therefore, a normal distribution was assumed. The leaching losses of the two teas were analyzed using a two-way (time vs. solution) ANOVA. Data were log-transformed prior to this analysis.

## Results and Discussion

### Effects of solute concentration on the teas’ leaching losses

We found that solute concentration, as well as soaking time, affected the leaching loss of both the green and rooibos teas (Fig. 1). The leaching loss of both teas was lower in tea solution compared to deionized water, and increased with increasing soaking time (Fig. 1a). This indicates that any leaching factor determined by submerging tea bags for several hours will be affected by solution concentration, including parameters such as dissolved organic carbon and salt concentrations. Therefore, it is difficult to determine an adequate leaching factor for the TBI method using a leaching experiment. We confirmed that our approach to determining leaching factors was appropriate, i.e., the use of multiple scenarios with different combinations, because leaching factors determined following several hours of submersion will likely change depending on solute concentration.

**Fig. 1.**
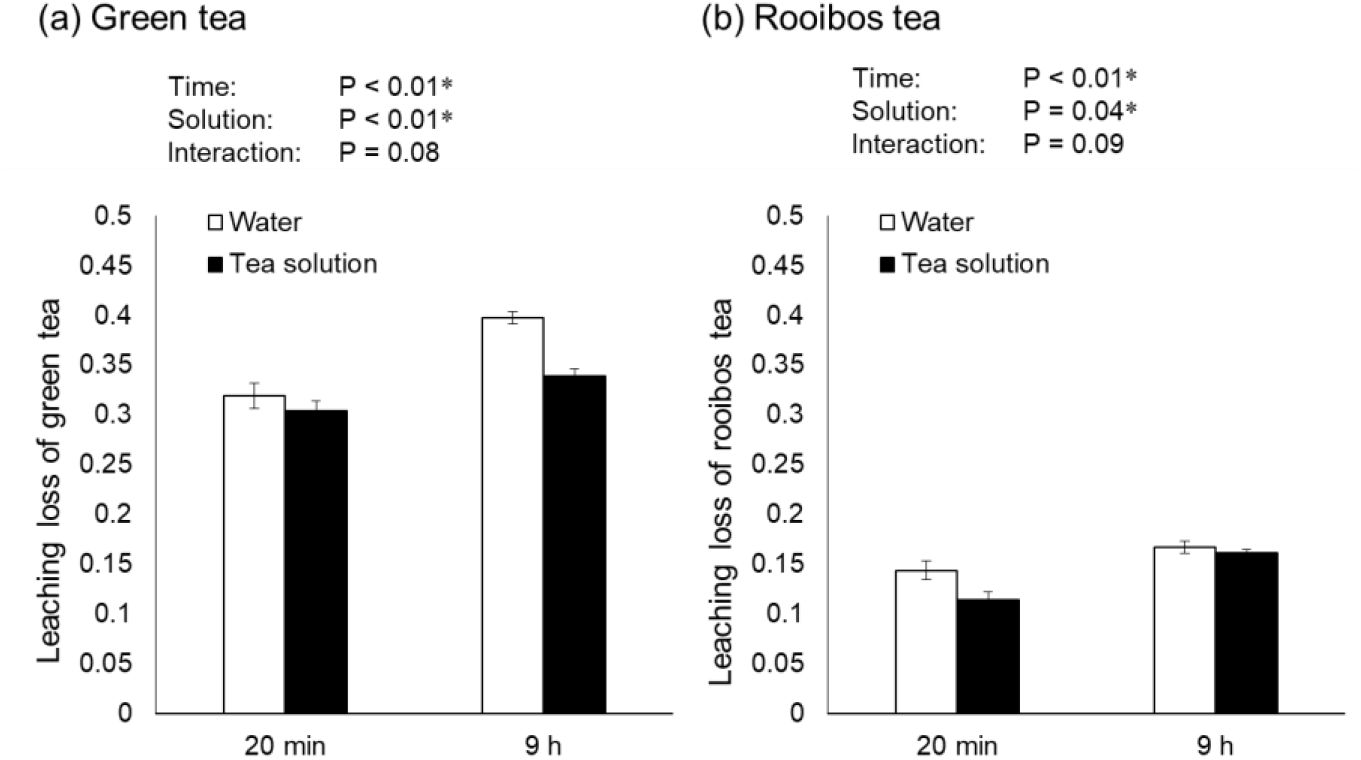
Relationships between solute concentration and the leaching ability of (a) green tea and (b) rooibos tea. Leaching losses were compared between deionized water and a tea solution for each tea type. Error bars indicate the standard error of three replicates.

### Evaluating the stabilization factor in aquatic environments

The remaining-mass ratios of the green tea samples at the end of the incubation period ranged from 0.3 to 0.6 (Fig. 2a–c), values within the range of those reported from aquatic environments (Peralta-Maraver et al., 2019; Seelen et al., 2019; Lalimi et al., 2018). The asymptote model did not fit the time-series green tea decomposition data well, because green tea decomposition did not plateau during the incubation period, at least not in the stream and pond water samples (Fig. 2a, b). Rather, green tea mass continued to decline until the end of the incubation period (Fig. 2). This result disagrees with those from a terrestrial system, where decomposition was found to plateau within the same period (Fig. 3a, Keuskamp et al. 2013). The continuous decline in mass therefore violates the plateau assumption of the TBI. This continuous decline was likely not due to slower overall decomposition rates in aquatic systems. In fact, the mass loss ratios of green tea in both the stream and pond samples were higher than those reported from terrestrial ecosystems (Djukic et al., 2018). We did not aim to investigate the potential causes of the differences in green tea decomposition curves between aquatic and terrestrial systems, and thus cannot speculate further on this point. However, we clearly demonstrated that the standard protocol for the TBI method is not suitable for determining *S* in aquatic environments. The inclusion of a leaching factor provides a negligible improvement in model fit, because it does not eliminate the issue of a decreasing trend in green tea mass loss (Fig. S1).

**Fig. 2.**
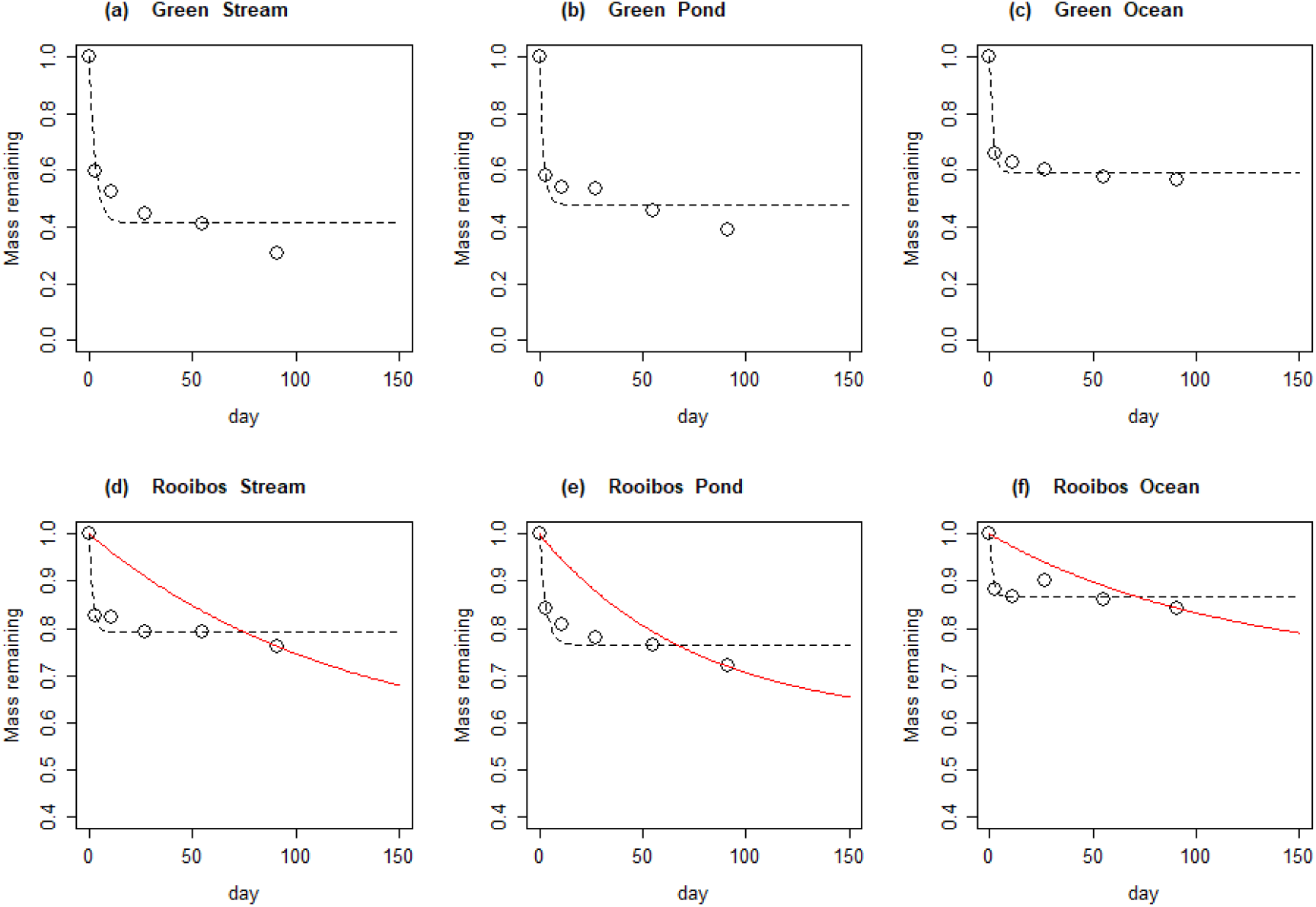
Relative remaining masses of green (a–c) and rooibos (d–f) tea bags in water samples taken from a stream (a, d), pond (b, e), and ocean (c, f) in Kumamoto, Japan. Open circles represent the average of four replicates. Tea bags were incubated in the dark and retrieved at 0, 3, 11, 27, 55, and 91 days after the start of the incubation. Dashed lines indicate the associated asymptote model: *W(t) = a * e*^*-kt*^ *+ (1-a)*, where *W(t)* is the mass remaining after incubation time *t, k* is a decomposition constant, *a* is the decomposable and *1-a* is the un-decomposable fraction of the teas. Solid red lines indicate the asymptote models describing rooibos tea decomposition, as determined by the TBI.

**Fig. 3.**
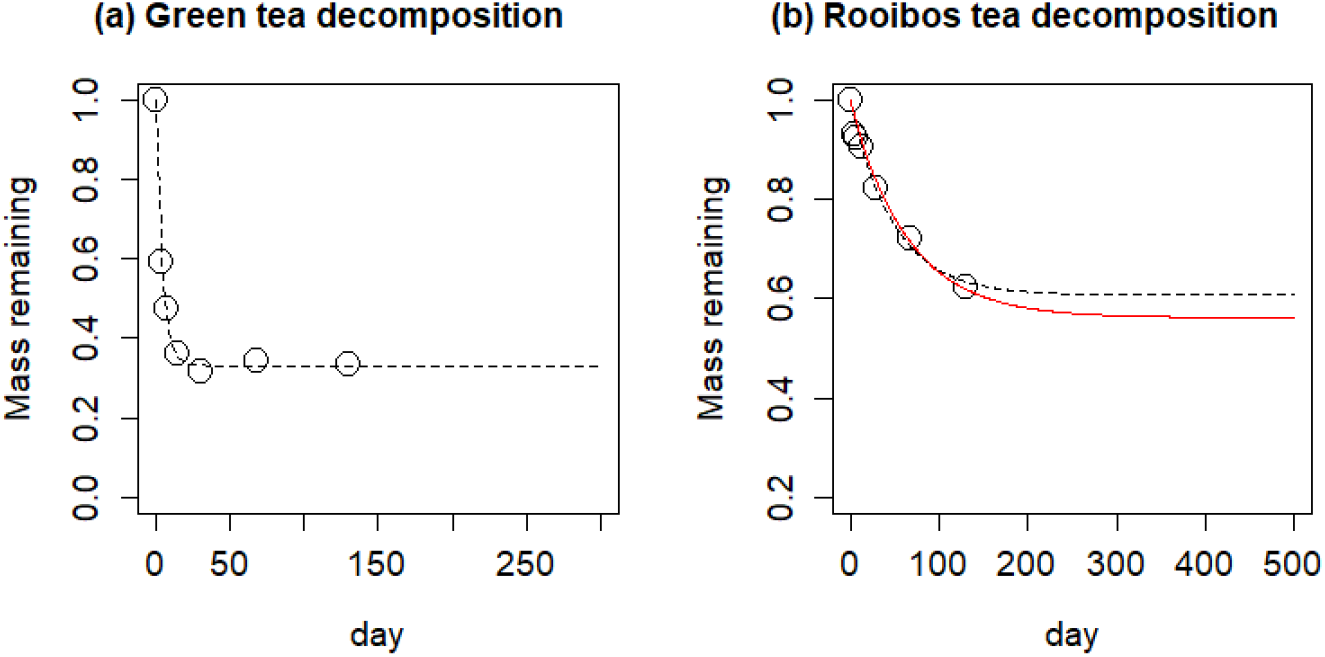
Relative remaining masses of (a) green and (b) rooibos teas in a terrestrial system. Data were obtained from Keuskamp et al. (2013) using Data Thief 3.0 (Tummers, 2006). Tea bags were incubated at 25°C in the dark for 0, 4, 7, 14, 30, 68, and 130 days. Dashed lines indicate asymptote models (reported by Keuskamp et al. 2013) see Fig. 2. The solid red line indicates the TBI-based model.

### Evaluating the decomposition constant and TBI-based asymptote models in aquatic environments

We found that *k* values determined using the TBI approach (0.0081, 0.013, and 0.0088 in stream, pond, and ocean samples, respectively) were at least one order of magnitude lower than those determined by fitting actual decomposition data (0.58, 0.33, and 0.72 in stream, pond, and ocean samples, respectively), indicating substantial disagreements between the TBI-based asymptote models and the models constructed using actual data (Fig. 2d–f). These results contrast with those obtained in a terrestrial ecosystem (Fig. 3b, Keuskamp et al. 2013), where these two models produced nearly identical results. We suggest that this discrepancy is the product of a violation of the assumption that the ratio of the decomposable to the total hydrolysable fraction is the same for rooibos and green tea. The decomposable hydrolysable fraction (*a*_*r*_) values determined by the TBI approach (0.46, 0.40, and 0.29 in stream, pond, and ocean samples, respectively) were much larger than those obtained from the actual data-based model (0.21, 0.24, and 0.13 in stream, pond, and ocean samples, respectively). The fact that green tea decomposition did not reach a plateau, such that *S* may have been overestimated, was not the reason for the discrepancy, because the overestimation of *S* should cause an underestimation of *a*_*r*_ (see eq. 2). Therefore, compensating for any overestimation of *S* would widen the differences between the TBI-based model and actual-data-based model. Subtracting the leachable fraction from the initial tea mass, as proposed by Seelen et al. (2019), improved the TBI-based asymptote model (Fig. 4). A larger leaching factor reduced the discrepancy between the TBI-based model and actual-data-based model for rooibos tea (Fig. 4). However, excluding some cases with ocean water samples, wherein the leaching factors were 80% and 100% of the maximum (Fig. 4k, l), the discrepancy remained. Therefore, the TBI approach may overestimate the hydrolysable fraction of rooibos tea, leading to an underestimation of *k* in asymptote models. We note that the asymptote model did not fit the real-world data well, and the discrepancy between the two model types may therefore have been overestimated. However, considering that we could not determine *S*, and that under- or over estimation of *S* leads to incorrect estimation of *k* (Mori et al., 2021b), we concluded that the standard protocol used for the TBI method, which was developed in a terrestrial ecosystem, is not applicable to aquatic ecosystems.

**Fig. 4.**
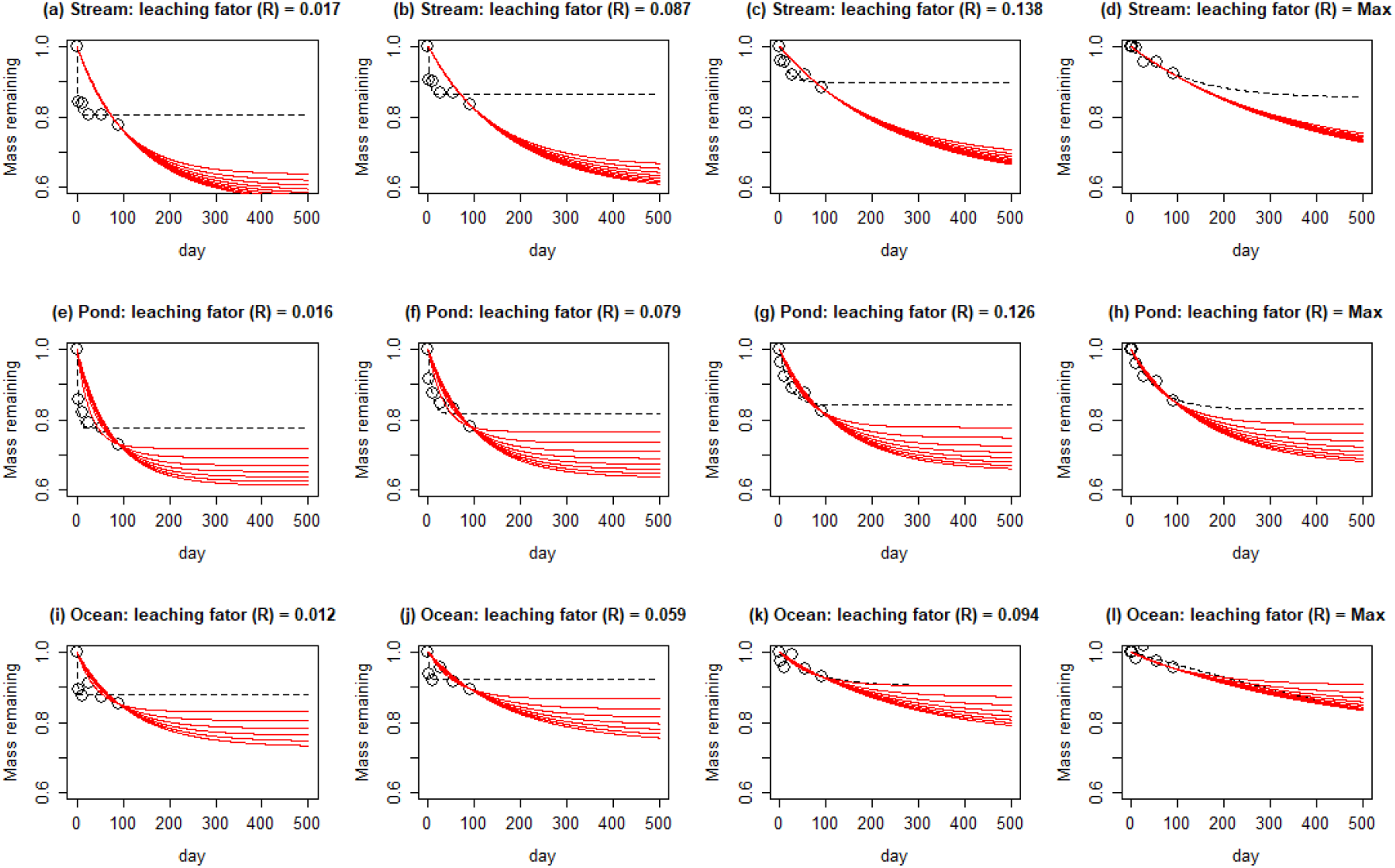
Leaching factor correction scenarios for the TBI-based asymptote model. Multiple scenarios with different leaching factors were tested. The remaining-mass values of rooibos tea at 0, 3, 11, 27, 55, and 91 days after the start of incubation are shown, assuming 10% (a, e, i), 50% (b, f, j), 80% (c, g, k), and 100% (d, h, l) of the maximum leaching factor of rooibos tea. Dashed lines indicate asymptote models. Solid red lines indicate TBI-based asymptote models, assuming 0%, 12.5%, 25%, 37.5%, 50%, 62.5%, 75%, 87.5%, and 100% of the maximum leaching factor of green tea. Water samples were obtained from a stream (a–d), pond (e–h), and ocean (i–l) in Kumamoto Prefecture, Japan. R indicates the leaching factor for rooibos tea.

### Future prospects for the use of the TBI method in aquatic environments

Although the TBI may be an unsuitable index in aquatic ecosystems, this does not necessarily negate the benefits of using tea bags as a standard material in assessments of litter decomposition rates in aquatic systems. For example, the remaining raw mass of both tea types could provide information on decomposition rates that is comparable on a global scale (Djukic et al., 2018). We found that the remaining mass of both teas following 91 days of incubation reflected the differences among the three water sample types used in this study (Fig. 5). This indicates that tea bags are sensitive enough to reflect differences in chemical composition and microbial community, both of which control litter decomposition (temperature was controlled in our study). Moreover, patterns in the remaining-mass ratios among the water sample types differed between the two tea types. The ratio was lowest in stream water for green tea and lowest in pond water for rooibos tea, although it was highest in ocean water for both types, likely because the high salt concentration inhibited decomposition (Contreras et al., 2017). Combining these two tea types could be advantageous for detecting differences in decomposition processes among different aquatic ecosystems.

**Fig. 5.**
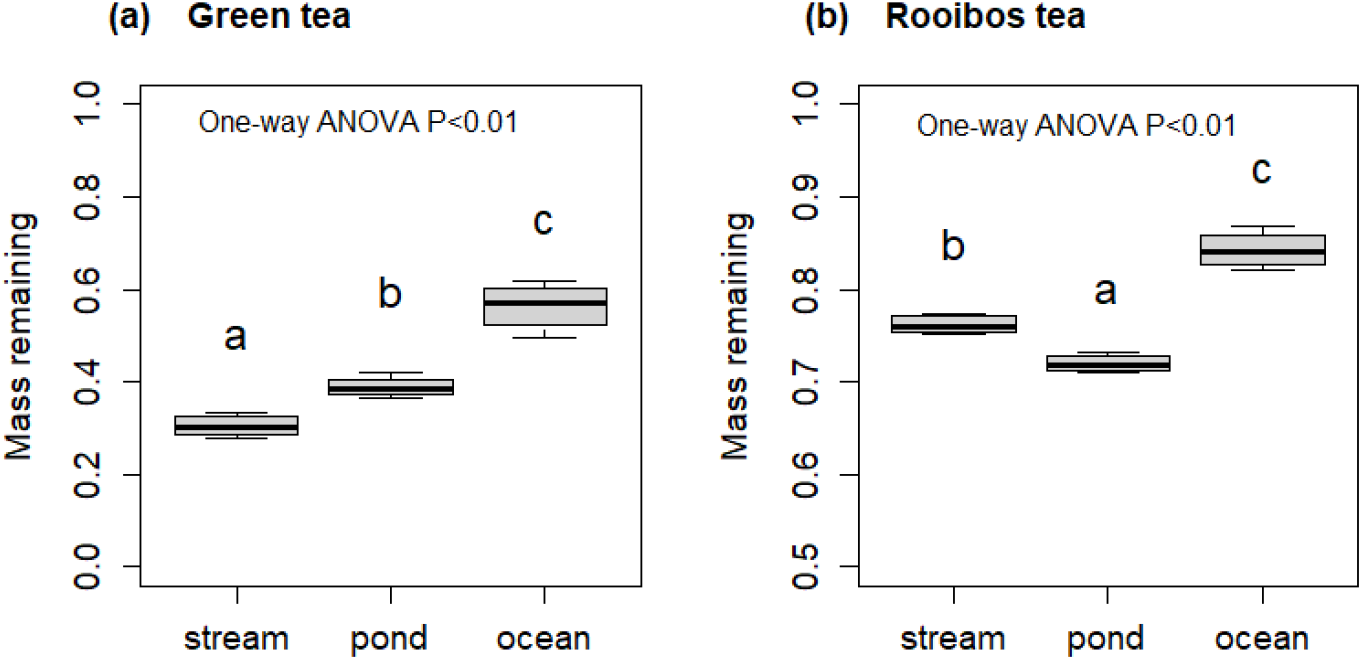
Boxplots representing the remaining mass values of (a) green and (b) rooibos tea bags at the end of a 91-day incubation period. Amounts relative to initial weights are shown. Letters indicate significant differences among groups, as determined by Tukey’s post hoc tests.

In addition, tea bags could still be used as standard materials to obtain time-series data and fit models in assessments of decomposition rates. In this scenario, an alternative model type could also be selected, as the asymptote model may not fit such data best. For example, we fitted a double exponential model, which is a generalized version of an asymptote model (i.e., an asymptote model is a special case of the double exponential model where *k2* = 0), and found improved fit relative to an asymptote model (Fig. 6). The AIC values resulting from double exponential models (−26.0, −28.3, −33.3, and −36.9 in green tea-stream, green tea-pond, rooibos tea-stream, and rooibos tea-pond, respectively) were lower than those from the asymptote model (−10.5, −12.9, −25.5, and −21.5 in green tea-stream, green tea-pond, rooibos tea-stream, and rooibos tea-pond, respectively). More complicated models, using more data with more frequent sampling and longer incubation periods, such as triple exponential models considering the decomposition of the acid-insoluble fraction, may provide important further information. This approach would negate a major advantage of the TBI method, i.e., only a single measurement is required to determine an asymptote model, but it is still beneficial because tea bags reduce the labor of preparing litter bags and the materials are highly standardized, which enables comparison between studies. In conclusion, we have demonstrated that the TBI approach is not applicable to aquatic environments, but we nevertheless suggest that tea bags are beneficial for assessing aquatic litter decomposition, so their potential application should be further assessed.

**Fig. 6.**
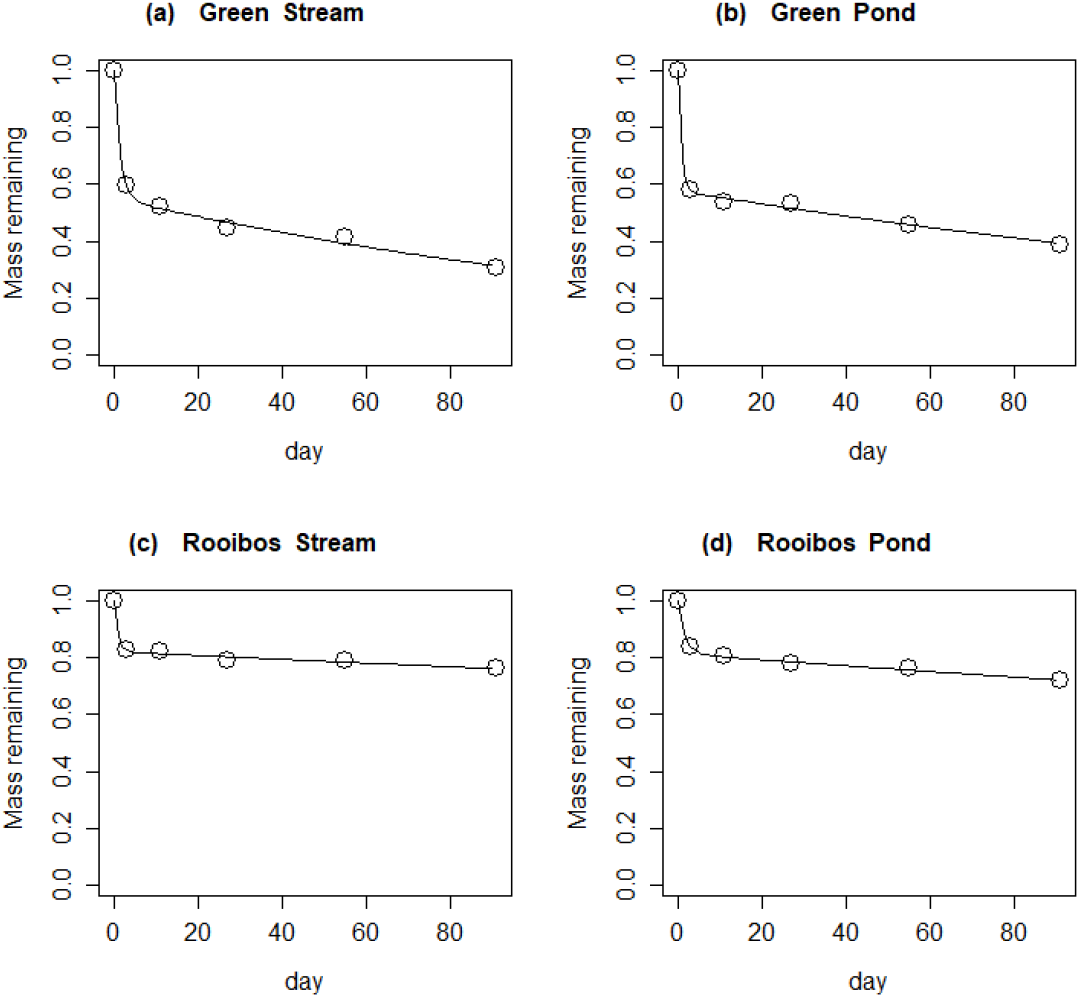
Time series data of green (a, b) and rooibos (c, d) teas in water samples taken from a stream (a, c) and pond (b, d) in Kumamoto, Japan. Solid lines indicate double exponential models: *W(t) = a * e*^*- k1t*^ *+ b * e*^*-k2t*^, where *W(t)* is the mass remaining after incubation time *t, k1* and *k2* are decomposition constants, and *a* and b represent organic matter fractions differing in decomposability. Each open circle represents the average of four replicates. Tea bags were incubated in the dark and retrieved at 0, 3, 11, 27, 55, and 91 days after the start of the incubation.

## Supplemental materials

**Fig. S1.**
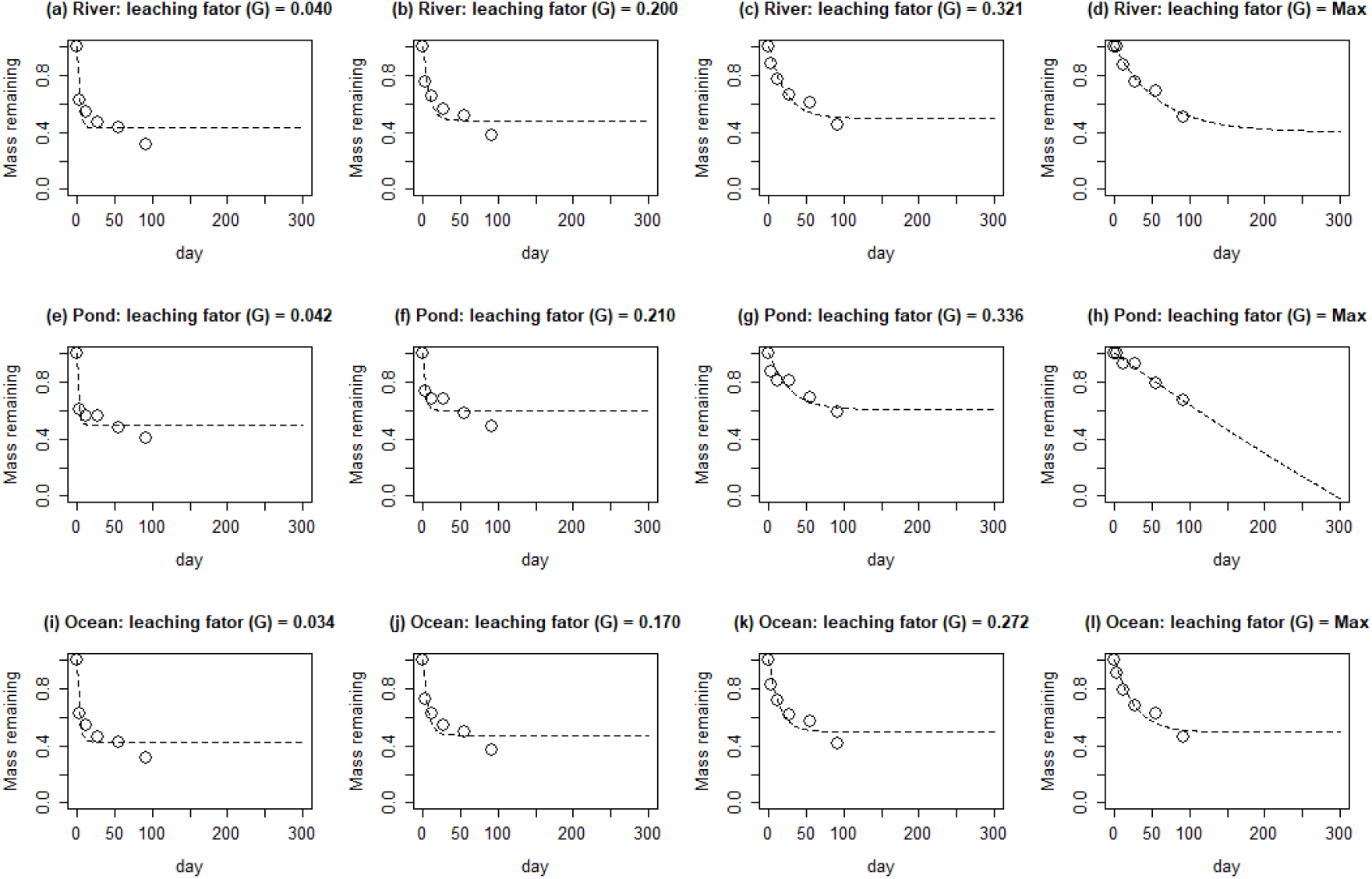
Leaching factor scenarios applied to time-series green tea decomposition data. The remaining-mass values of green tea at 0, 3, 11, 27, 55, and 91 days after the start of the incubation are shown, assuming 10% (a, e, i), 50% (b, f, j), 80% (c, g, k), and 100% (d, h, l) of the maximum leaching factor of green tea. Dashed lines indicate asymptote models. Data were obtained from a stream (a–d), pond (e–h), and ocean (i–l) in Kumamoto Prefecture, Japan. G indicates the leaching factor of green tea.

